# Evolutionary and functional connections between the typhoid toxins encoded by *S.* Typhi and *S. bongori*

**DOI:** 10.1101/2024.05.21.595087

**Authors:** Antonio J. Chemello, Casey C. Fowler

## Abstract

The virulence properties of numerous important bacterial pathogens have been shaped by the horizontal acquisition of an AB5-type toxin. A major obstacle to the fruitful horizontal transfer of an AB5-type toxin amongst bacteria that infect divergent host species is that these toxins bind specific glycan receptors on target host cells to mediate toxin uptake, and glycosylation patterns vary substantially between different host species. In this study, we explore the evolutionary and functional connections between the related typhoid toxins that are encoded by *S. enterica* serovar Typhi, a human-adapted pathogen, and by *S. bongori*, which infects cold-blooded animals. We uncover genetic fingerprints that indicate *S*. Typhi’s typhoid toxin islet evolved from a larger virulence islet similar to *S. bongori*’s, and show that *S*. Typhi’s glycan-binding PltB subunit is more similar to *S. bongori* PltB than it is to many diverse PltB sequence variants found in *S. enterica*. We further show that, surprisingly, *S*. Typhi strains that encode the *S. bongori* typhoid toxin islet intoxicate infected human epithelial cells with substantially greater potency than wild-type *S*. Typhi. The results of this study therefore suggest that the unexpected potency of the *S. bongori* typhoid toxin toward mammalian cells paved the way for sequence variants derived from this species to be acquired by - and effectively integrated into the virulence program of - the human-adapted pathogen *S*. Typhi.

## INTRODUCTION

Typhoid toxin is produced by the human pathogen *Salmonella enterica* serovar Typhi (*S*. Typhi), the cause of typhoid fever [1,2]. It’s an A_2_B_5_ toxin composed of two active (A) subunits, PltA and CdtB, and a delivery (B) subunit homopentamer composed of PltB, or an alternate B subunit, PltC [3–5]. Typhoid toxin is expressed by *S*. Typhi within the intracellular niche adopted following host cell invasion, the *Salmonella* containing vacuole (SCV) [6]. Toxin is then secreted, exocytosed from the cell, and then binds receptors on target host cells, leading to cellular intoxication [3,7,8]. Trafficking steps are mediated by the toxin’s B subunits through engaging specific sialoglycan receptors [7–9]. Typhoid toxin’s known biological effects stem from CdtB, which is a DNAse that causes DNA damage that provokes G2/M cell cycle arrest and cellular senescence [10]. Typhoid toxin is a key virulence factor that plays a role in disease progression and likely contributes to *S*. Typhi’s ability to establish persistent infections [4,11,12].

*Salmonella* is composed of two species, *S. enterica* and *S. bongori. S. bongori* and five of the six *S. enterica* subspecies primarily infect cold-blooded animals, but the *enterica* subspecies is adapted to warm-blooded animals and causes most human infections (Fig 1A) [13,14]. Typhoid toxin is absent from most subsp. *enterica* serovars, including those that most commonly cause gastroenteritis (e.g. Typhimurium), but genetically similar typhoid toxin islets are found at the same genomic location in the typhoidal serovars and a distinct clade of the subspecies (“clade B”) [5,15–18]. Prophage-encoded typhoid toxin loci are also found in individual strains of various subsp. *enterica* serovars that generally lack typhoid toxin [5,16].

**Fig 1:**
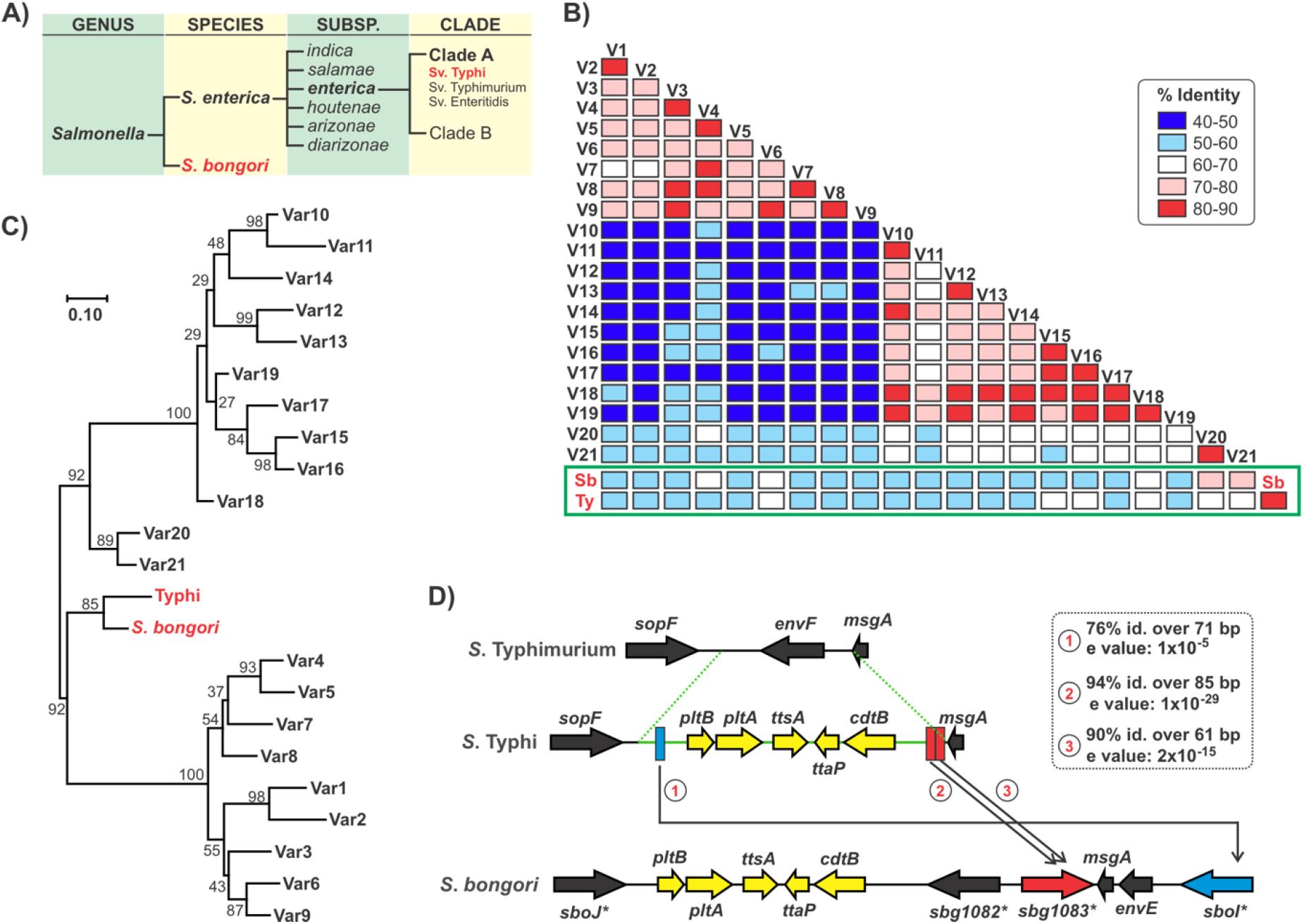
Sequence analyses identify evolutionary connections between the *S. bongori* and *S*. Typhi typhoid toxin loci. **(A)** Streamlined overview of *Salmonella* taxonomy highlighting the lineages most relevant to this study. **(B)** Percent sequence identity matrix for all pairwise combinations of representative PltB variants. 23 representative PltB sequences, each of which is <90% identical to all others, were identified in the NCBI nr protein database and aligned using Clustal Omega. The close sequence relationship between the *S. bongori* (Sb) and *S*. Typhi (Ty) variants is highlighted using a green box. PltB sequences and alignments are shown in Fig S1-S2. **(C)** The PltB variants described in (B) were used to generate a phylogenetic tree using the MEGA (Molecular Evolutionary Genetics Analysis) V11 software using the maximum likelihood method and the WAG substitution model. A bootstrap method with 300 total replicates was used, and the numbers at the nodes represent the bootstrap support values. **(D)** BLASTn search of the *S*. Typhi typhoid toxin islet (the boundaries of which were identified via comparisons to the corresponding locus in *S*. Typhimurium, green dashes) against *S. bongori* revealed three segments of DNA sequence similarity between the noncoding segments that flank the *S*. Typhi typhoid toxin and genes adjacent to the typhoid toxin locus on the larger *S. bongori* virulence islet. The three segments are shown as rectangles color-coded to correspond to the relevant *S. bongori* gene. Sequence alignments for these regions are shown in Fig S3. Analysis of the distributions of these *sbg1082*/*sboI* sequence similarity patches amongst other typhoid toxin-encoding lineages is shown in Fig S4. Asterisks denote large genes that are not shown to scale in order to maintain visual clarity.

This sporadic distribution suggests typhoid toxin-encoding subsp. *enterica* lineages acquired this toxin horizontally. Typhoid toxin is only found in *Salmonella*, but is broadly distributed in lineages outside of subsp. *enterica* [5,18]. Interestingly, *S*. Typhi typhoid toxin is more genetically similar to *S. bongori* typhoid toxin than to typhoid toxins encoded by other *S. enterica* subspecies, and *S. bongori* typhoid toxin intoxicates human cells during infection [18]. This prompted us to explore the connections between the typhoid toxins encoded by *S*. Typhi and *S. bongori*.

## RESULTS

### Genetic connections between the *S*. Typhi and *S. bongori* typhoid toxin loci

For many AB_5_-type toxins, including typhoid toxin, the predominant functional differences between sequence variants track to the delivery subunits [19,20]. Glycosylation patterns can vary substantially between hosts, and evolving to target receptors in one species can lead to reduced potency against another [21–23]. To explore the relationship between the *S. bongori* and *S*. Typhi typhoid toxins, we therefore compared their PltB subunits in the broader context of the PltB diversity present in *Salmonella*. We queried the NCBI nr protein database and identified 23 representative PltB variants, such that each representative is <90% identical to the others, and all PltBs identified grouped with (were >90% identical to) a representative. Most PltBs entries in subsp. *enterica* group with *S*. Typhi PltB, including those in the typhoidal and clade B serovars, whereas all PltBs that group with the *S. bongori* representative appear to be from *S. bongori* (Table S1). The remaining 21 PltB variants are scattered around *S. enterica*, with multiple different variants found in subsp. *diarizonae, arizonae, houtenae* and *salamae*.

Interestingly, most variants are found in certain strains of various subsp. *enterica* serovars, consistent with the observation that prophage-encoded typhoid toxins are widespread and genetically diverse [16]. We aligned all 23 PltB representative sequences and generated a percent identity matrix and a phylogenetic tree (Fig 1B-C, S1, S2). The average sequence identity across all pairwise combinations of variants was 60.8%, highlighting the striking PltB sequence diversity in *Salmonella*. Importantly, *S*. Typhi and *S. bongori* PltBs exhibited much higher levels of sequence identity to one another (84%) than to any other representative (50-72%). This is also reflected in the phylogenetic tree, which predicts that *S*. Typhi and *S. bongori* PltB are closely related and share a unique common ancestor. This analysis revealed that *S*.

Typhi PltB is more similar to *S. bongori* PltB than it is to many diverse PltB variants that are scattered across the *S. enterica* lineage.

We next mined NCBI DNA sequence databases, identified 78 total *S. bongori* genomes, and found that each encodes a typhoid toxin locus that is >98% identical to that of *S. bongori* SARC 11 (Supplemental Datafile 1). This indicates that the typhoid toxin locus is part of the *S. bongori* core genome and is thus likely of ancient origins in this species. In *S. bongori*, the typhoid toxin locus is part of a larger *S. bongori*-specific virulence island that encodes a serine protease autotransporter (*sbg1083*) and two type III secretion system effector proteins, *sboI* and *sboJ* (Fig 1C) [24]. Importantly, a BLAST search of the full *S*. Typhi islet, which includes ∼750 bp noncoding sequence on either side of the coding genes, revealed that short patches of these noncoding flanks exhibit significant sequence similarity to the *sbg1083* and *sboI* genes from the larger *S. bongori* islet (Fig 1D, S3). These evolutionary fingerprints indicate that *S*. Typhi typhoid toxin evolved from a toxin that was part of a larger *S. bongori*-like virulence locus. These homology regions are also found in subsp. *enterica* clade B and certain prophage-encoded typhoid toxin loci that are genetically similar to the *S*. Typhi islet, but are absent from genetically divergent typhoid toxin loci, including those in other *S. enterica* subspecies (Fig S4).

Collectively, these data suggest that the *S. bongori* typhoid toxin locus was ancestral to the locus found in the vast majority of typhoid toxin-encoding strains associated with human infection, including the typhoidal serovars.

### *S*. Typhi strains that encode the *S. bongori* typhoid toxin more potently intoxicate human cells than WT S. Typhi

To investigate the functional relationship between the *S. bongori* and *S*. Typhi typhoid toxins, we devised an experiment to mimic the horizontal transfer of the *S. bongori* typhoid toxin islet into *S*. Typhi. We first generated a *S*. Typhi strain featuring a deletion of all typhoid toxin genes (ΔTT), and a plasmid carrying the *S. bongori* typhoid toxin locus on a mariner transposon (Fig 2A). We then isolated 12 transposon mutants wherein the *S. bongori* typhoid toxin islet randomly inserted into the *S*. Typhi genome; the insertion sites of 4 representative mutants were mapped, revealing unique locations (Fig S5). All mutants were used to infect human epithelial cells, and we assayed for CdtB-mediated intoxication using cell cycle analysis (Fig 2B).

**Fig 2:**
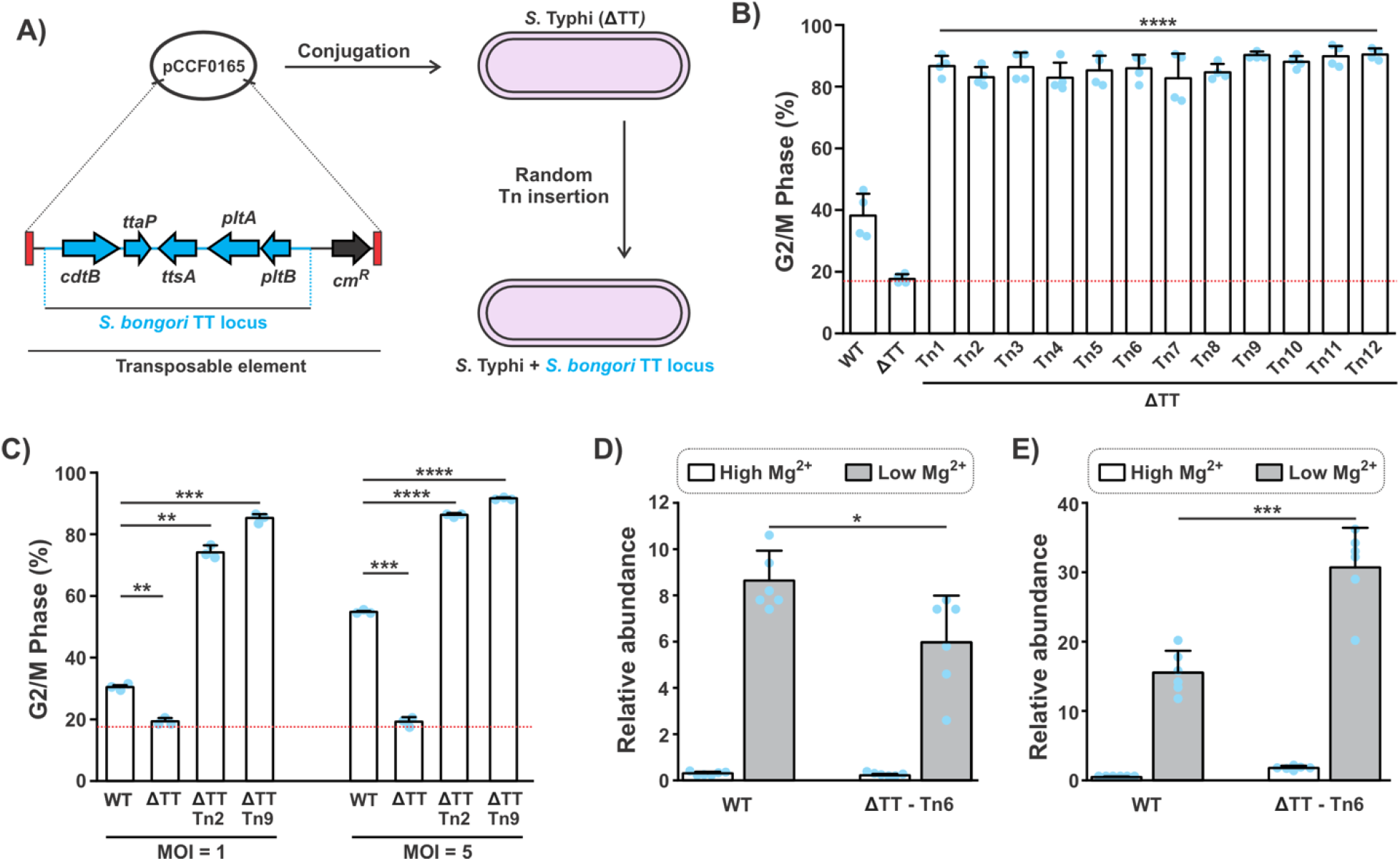
*S*. Typhi strains that encode the *S. bongori* typhoid toxin more potently intoxicate human epithelial cells during infection than WT *S*. Typhi. **(A)** Schematic outlining the generation *S*. Typhi strains carrying the S. bongori typhoid toxin locus. The *S. bongori* typhoid toxin, including 500 bp on either side of the terminal coding genes to capture the native promoter sequences, was cloned into the transposable element of a mariner transposon plasmid. Conjugative transfer of this suicide vector into a *S*. Typhi strain carrying deletions to the complete typhoid toxin islet as well as to *pltC* (which encodes an alternative typhoid toxin delivery subunit) yielded strains wherein *S. bongori*’s typhoid toxin island was randomly inserted into the *S*. Typhi genome. **(B and C)** *S*. Typhi strains, including WT, a typhoid toxin deletion strain (ΔTT), and transposon mutant strains (Tn) wherein the *S. bongori* typhoid toxin was introduced into the ΔTT strain, were used to infect HeLa cells at an MOI of 3 for (B), or as indicated in (C). At 72 hours post infection, cells were collected, fixed, strained using propidium iodide (PI), and analyzed by flow cytometry-based cell cycle analysis to determine the levels of CdtB-induced G2/M cell cycle arrest. Bars represent the average percentage of cells in the G2/M phase, error bars represent the standard deviation, and the red dotted line shows the average G2/M levels in unintoxicated control cells. Tukey’s test was used to determine statistical significance [compared to WT for (B), or as indicated in (C)]. **** P < 0.0001, *** P < 0.001, ** P < 0.01. **(D and E)** RT-qPCR analysis of transcript levels of *pltA* (D) and *cdtB* (E) in WT *S*. Typhi compared to a representative transposon mutant encoding the *S. bongori* typhoid toxin islet. Strains were grown for 6 hours in either low Mg^2+^ (10 μM) medium that stimulates typhoid toxin expression, or a high Mg^2+^ (1 mM) medium that does not, and RNA was extracted and analyzed by RT-qPCR. Transcript levels were first calibrated using genomic DNA; *pltA* and *cdtB* levels were then normalized to the housekeeping gene *dsbC*. Bars represent average relative transcript levels and error bars show standard deviations. Statistical significance was assessed using two tailed t-tests of the indicated comparisons: *** P < 0.001, * P < 0.05.

Surprisingly, we found that all 12 mutants elicited significantly greater cellular intoxication than the *S*. Typhi control strain encoding the WT typhoid toxin islet (WT). Indeed, mutants carrying the *S. bongori* toxin intoxicated more potently at a multiplicity of infection (MOI) of 1 than the WT *S*. Typhi strain did at an MOI of 5 (Fig 2C). Modest (<2-fold) differences in typhoid toxin gene expression between strains carrying the *S*. Typhi or *S. bongori* typhoid toxins suggest this is not the cause of differential toxicity (Fig 2D-E). Collectively, these results show that, in *S*. Typhi, the *S. bongori* typhoid toxin more potently intoxicates human cells than its native typhoid toxin. This is congruent with the evolutionary connections noted above, and provides a functional explanation for why *S. bongori*-derived typhoid toxins are predominant amongst *S. enterica* lineages associated with mammalian hosts.

## DISCUSSION

This study presents findings that support the premise that *S. bongori* typhoid toxin was ancestral to that of *S*. Typhi and other subsp. *enterica* lineages of clinincal relevance. It is unlikely that *S*. Typhi inherited typhoid toxin directly from *S. bongori*; rather, this was presumably a multi-step process that was mediated by phages. Indeed, there are strains featuring typhoid toxin-encoding prophages at the locus where typhoid toxin is found in *S*. Typhi [16]. One possibility is that *S. bongori* typhoid toxin was acquired by phages and was subsequently transferred into an early ancestor of subsp. *enterica* clade B. *S*. Typhi, which emerged ∼10-43 thousand years ago from a (“clade A”) ancestor that presumably lacked typhoid toxin [25], may have acquired typhoid toxin from a clade B strain, either directly (e.g. during co-infection of a host) or indirectly, via a phage intermediate.

AB5-type toxins are typically genetically mobile, and can be transformative virulence factors. A major barrier to a newly-acquired toxin’s capacity to benefit a bacterium is that it must effectively engage glycan receptors on target host cells, which can be highly variable amongst different species and cell types [19,23]. *S. bongori* PltB’s capacity to effectively engage mammalian receptors was presumably a major factor underlying *S. bongori*-like typhoid toxin variants becoming dominant in subsp. *enterica*, rather than the many other sequence variants present in *S. enterica*. Previous studies have highlighted the adaptation of typhoid toxin to *S*. Typhi’s unique pathogenesis and to the human host [8,9,26]. *S*. Typhi PltB specifically engages Neu5Ac-terminated sialoglycans found on human receptors (humans lack the enzyme that converts Neu5ac to Neu5Gc in other animals), and PltB’s intracellular trafficking during infection appears to be adapted to *S*. Typhi and is less efficient in nontyphoidal serovars [8,9]. Given the remarkable potency of the *S. bongori* typhoid toxin when expressed in *S*. Typhi, it is likely that *S*. Typhi fortuitously acquired a toxin that was well-adapted to a human host that initially evolved in a pathogen that is not associated with mammals.

## METHODS

### *In silico* typhoid toxin analyses

Representative PltB variants were identified via iterative BLASTp searches as described in Supplemental Methods. Sequences were aligned and percent identity matrices were generated using Clustal Omega, and a phylogenetic tree was generated by MEGA V11.0.13. The set of sequenced *S. bongori* genomes in the NCBI nr/wgs DNA sequence databases, and which encode typhoid toxin, was determined using BLASTn searches restricted to the *S. bongori* taxid (54736) as described in Supplemental Methods. Typhoid toxin loci with regions of sequence similarity to *sbg1083* and *sboI* were identified using targeted BLASTn searches against typhoid toxin encoding lineages, followed by manual investigations into the compositions and genome contexts of numerous typhoid toxin islets. Further details available in Supplemental Methods.

### Analysis of *S*. Typhi strains encoding *S. bongori* typhoid toxin

The *S. bongori* typhoid toxin locus was cloned into the transposable element of pSB4807 [6] and conjugated into CCF0175, a *S*. Typhi ISP2825 derivative featuring deletions of the typhoid toxin islet and *pltC* as described [6]. Transposon mapping used Sau3AI digests of genomic DNA to enable adaptor ligation; adaptors facilitated a two-PCR amplification of the transposon insertion region, which was sequenced. RNA extraction and RT-qPCR was conducted as described following growth in TTIM [18]; *pltA* and *cdtB* levels were calibrated using standard curves for genomic DNA and normalized to *dsbC* levels. For intoxications assays, cell cycle analysis was performed on infected HeLa cells using flow cytometry-based analysis of PI-strained cells as previously described [18]. Further details available in Supplemental Methods.

## SUPPLEMENTAL INFORMATION

Supplemental information includes Supplemental methods, 5 figures, 1 table, and 1 supplemental dataset (Excel file).

## AUTHOR CONTRIBUTIONS

C.C.F. performed some *in silico* analyses, A.J.C. performed some *in silico* analyses and all wet lab experiments. C.C.F. and A.J.C. analyzed data and contributed to experimental design. C.C.F. and A.J.C. wrote the paper.

## ACKNOWLEDGEMENTS

This work was supported by NSREC discovery grant (Grant number: RGPIN-2020-03964 to C.C.F.) and a start-up grant provided by the University of Alberta Faculty of Science (to C.C.F).

A.J.C. was supported in part by an Alberta Innovates Summer Studentship award and an NSREC Undergraduate Student Research Award (NSERC-USRA). Flow Cytometry was performed at the University of Alberta, Faculty of Medicine and Dentistry Flow Cytometry Facility, which received financial support from the Faculty of Medicine and Dentistry and the Canadian Foundation for Innovation (CFI) awards to contributing investigators.

